# A reference-free algorithm discovers regulation in the plant transcriptome

**DOI:** 10.1101/2024.05.23.595613

**Authors:** Elisabeth Meyer, Evan V. Saldivar, Marek Kokot, Bo Xue, Sebastian Deorowicz, Seung Y. Rhee, Julia Salzman

## Abstract

Most plant genomes and their regulation remain unknown. We used SPLASH - a new, reference-genome free sequence variation detection algorithm - to analyze transcriptional and post-transcriptional regulation from RNA-seq data. We discovered differential homolog expression during maize pollen development, and imbibition-dependent cryptic splicing in Arabidopsis seeds. SPLASH enables discovery of novel regulatory mechanisms, including differential regulation of genes from hybrid parental haplotypes, without the use of alignment to a reference genome.

## Main text

The study of plant genomes and transcriptomes is fundamental to advancing basic biological science, crop resilience, and ecosystem stewardship. Today, virtually all plant genomic analysis begins with alignment to a reference genome or transcriptome. However, alignment-based approaches are limited by how well aligners perform and how well the available reference genome approximates the true genome. The assembly of plant genomes is particularly challenging due to complexities such as its intrinsic plasticity, high fractions of repetitive sequence^1,2^, polyploidy, and gene duplications^3,4^.

These problems, while acute in plants, are a general obstacle to discovery across the tree of life. To address them, we recently introduced a new approach to analyze regulation of genomes and transcriptomes using an ultra-efficient, reference-free, statistical approach called SPLASH^7,8^. In brief, SPLASH identifies statistically significant sample-specific sequence variation directly from raw reads, bypassing alignment to a reference^7,8^. SPLASH processes raw FASTQ reads and records sequences of a set length (i.e. kmers) called “anchors” within the reads that flank variable kmers. SPLASH then identifies anchors whose relative abundance depends on the experimental sample; these anchors are deemed significant. SPLASH can detect numerous biological processes that diversify sequence, including alternative splicing and differential homolog expression (Supplementary Fig. S1A; Methods). SPLASH can also identify sequence variation in organisms co-associating with the host. The significant anchors and their associated variants can be used to investigate the mechanism behind the variation (Supplementary Fig. S1B).

To illustrate the power of SPLASH to discover (post-)transcriptional regulation in plants, we re-analyzed four RNAseq datasets from maize (*Zea mays), Arabidopsis thaliana*, and sorghum *(Sorghum bicolor)* (Fig. 1, Supplementary Information). SPLASH makes thousands of discoveries in each dataset (Supplementary Information) without information about sample identity or reference genomes (Methods).

**Figure 1.**
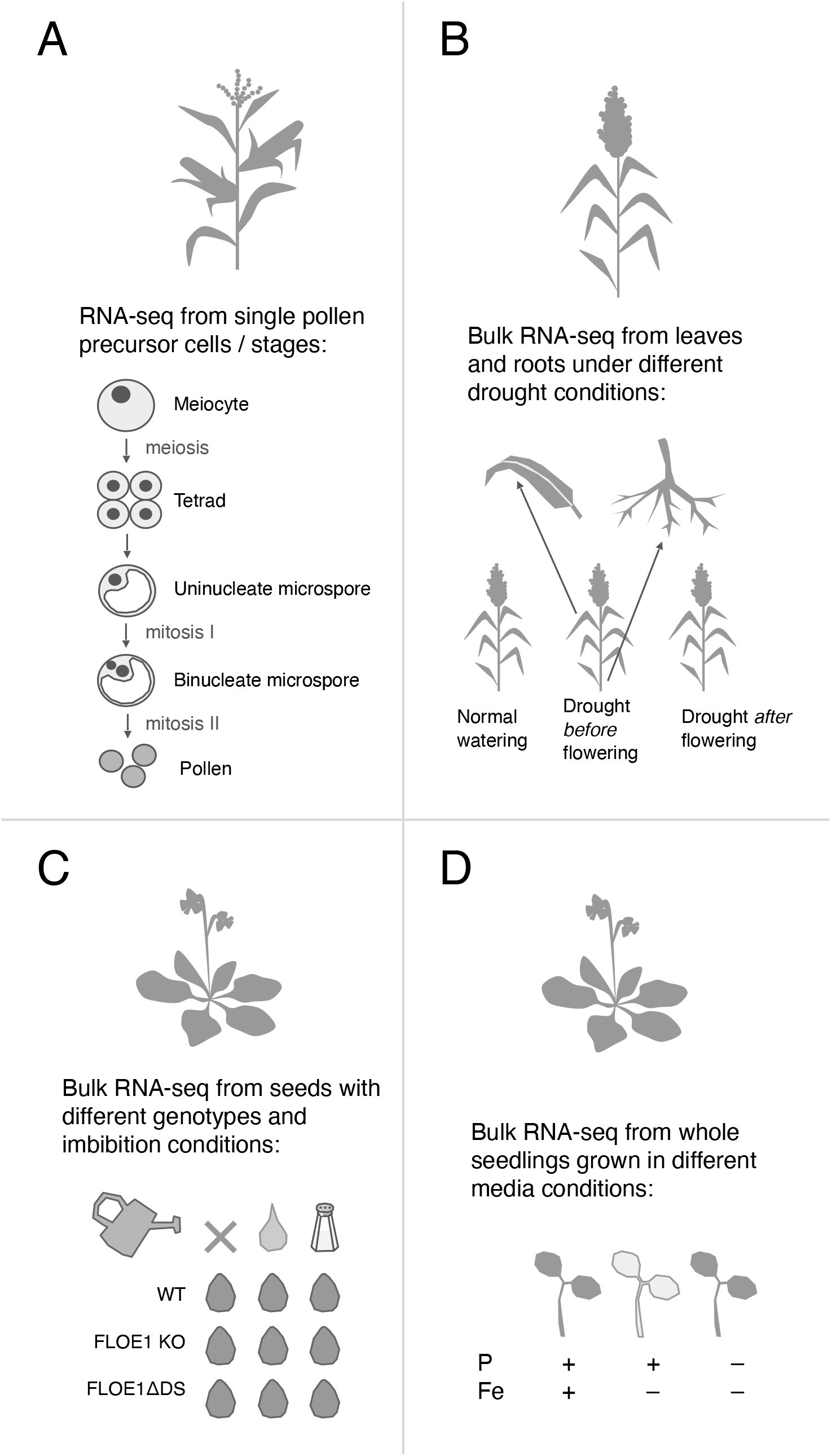
Overview of datasets analyzed with SPLASH in this paper. A: a single-cell study of maize pollen precursors. B: A bulk RNA-seq study of sorghum under different drought conditions. C: A bulk RNA-seq study of *Arabidopsis* seed germination, with wild-type and mutant strains, in conditions of no water, plain water imbibition, or salt water imbibition. D: A bulk RNA-seq study of *Arabidopsis* seedlings under iron or iron/phosphorus deficiency.

### SPLASH discovers complex transcript abundance patterns associated with plant genes

We re-analyzed a single-cell study of maize pollen precursors derived from a hybrid line between the nested association mapping (NAM) parental lines B73 and A188^10^. In this dataset^10^, there were 8,989,404 total significant anchors. SPLASH discovered developmental regulation of genes across both parental alleles, without requiring alignment to a reference genome (Fig. 2A, SFig. 2A). For example, the B73 allele of Zm00001eb173470 (preliminary annotation ID: Zm00001e021816), which encodes a ribosome-like protein, is expressed more highly over the A188 allele during meiosis. However, the A188 allele is more dominantly expressed than the B73 allele in other stages of pollen precursor development such as prophase I, mitosis I, and mitosis II. The complex developmental regulation of this gene, found by SPLASH without using either of the reference genomes, underscores the simplicity of a reference-free approach for discovery in plants derived from hybridization.

**Figure 2.**
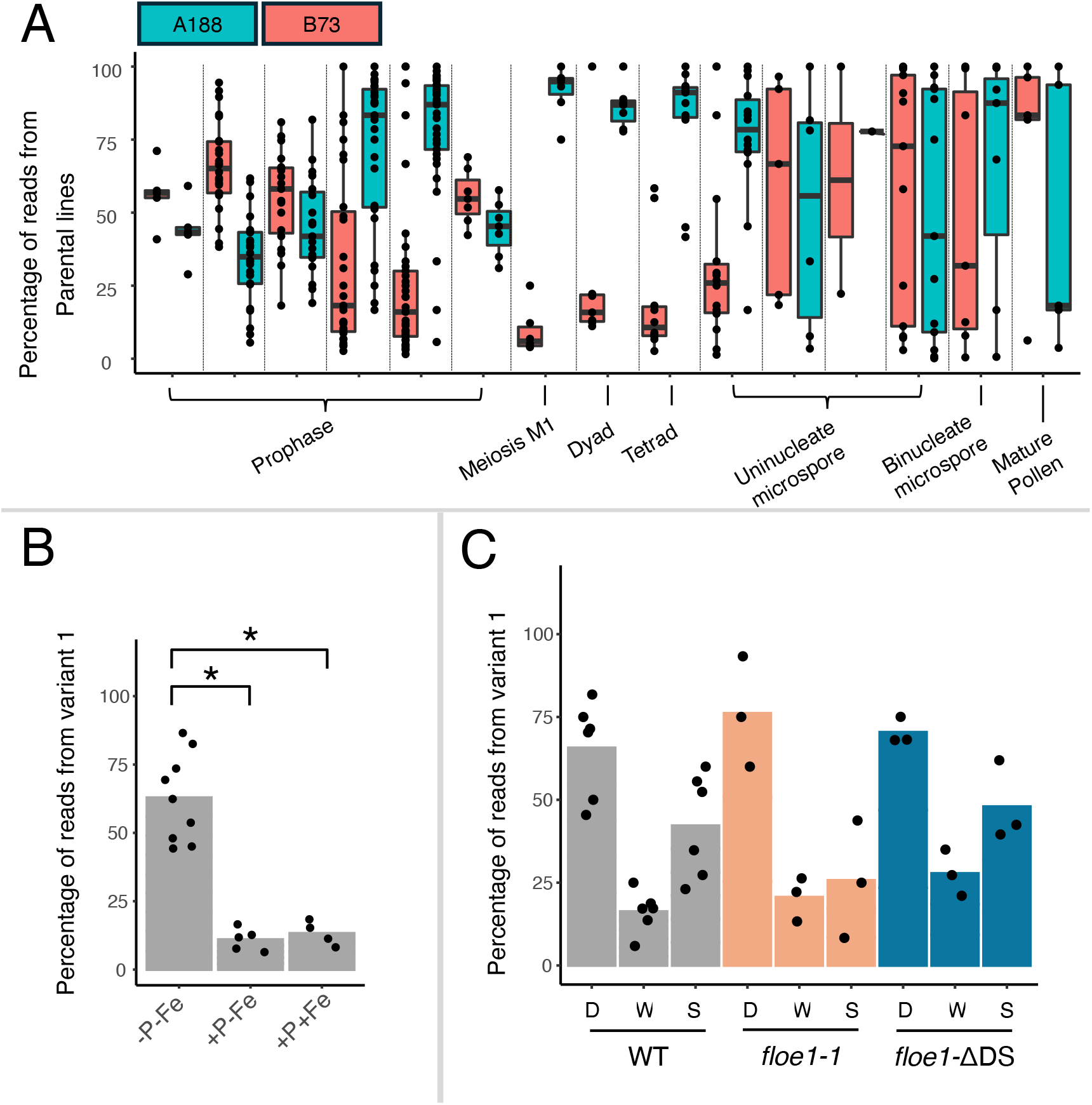
Percentage of reads within each sample from each of the top two variants, calculated as number of reads from that particular variant divided by total number of reads from the anchor. Bar height represents mean; individual datapoints are superimposed. Whether variation depends on experimental condition was tested in two different ways. For A and C, we used a generalized linear model to test whether the experimental condition/developmental stage could be predicted from the proportion of variants (Methods). For B, pairs of conditions were compared using a two-proportions z-test; asterisks indicate that the proportion of variant 1 reads is significantly different (p-value < 0.05) between experimental conditions (Methods). For the *Arabidopsis* FLOE1 dataset, only wild type (WT) samples were tested for significance. A: Maize pollen dataset: the top two variants for this anchor align to alleles of Zm00001eb173470 from B73 and A188. The differential expression of these alleles varies by pollen stage. Note: this anchor was only found in 260 out of 642 samples. B: *Arabidopsis* iron/phosphorus deprivation dataset: variant 1 maps to a splice junction in AT1G74270 (ribosomal protein L35Ae), while variant 2 includes the intron. -P-Fe indicates the phosphorus and iron doubly deprived condition; +P-Fe is only iron deprivation; and +P+Fe indicates no deprivation. C: *Arabidopsis* FLOE1 dataset: variant 1 maps to an annotated splice junction between exons in AT2G36720, but variant 2 maps to a cryptic splicing event from inside an intron to an exon. WT (wild type), *floe1-1* (FLOE1 deletion mutant), and *floe1-ΔDS* (FLOE1 mutant with disordered region deleted) indicate the different seed genotypes; D (dry), W (wet), and S (salt) indicate the imbibition conditions for the seeds.

In addition, we re-analyzed a study of wild type *Arabidopsis* seedlings, which showed that chlorosis (loss of chlorophyll) induced by iron deficiency involves a phosphorus-dependent pathway, so iron/phosphorus doubly-deprived plants have a “stay-green” phenotype^12^. However, the genetic pathway that underpins this phenotype is not fully elucidated, warranting further study. A snapshot of SPLASH’s findings includes condition-dependent alternative splicing in gene AT1G74270 (ribosomal protein L35Ae): the phosphorus-deficient samples do not express the spliced isoform, whereas the samples not deprived of phosphorus express the spliced version (Fig. 2B, SFig. 2B). To our knowledge, this is the first example of a splicing regulation in a plant ribosomal protein gene impacted by nutrient deprivation.

Finally, we re-analyzed a study of *Arabidopsis* that discovered the gene FLOE1 as a regulator of seed germination^14^. This study showed that FLOE1 controls seed germination under water-limiting conditions and senses water availability through condensate formation. Condensate formation is important for FLOE1’s ability to control germination. How FLOE1’s condensate formation controls germination remains unknown. The study includes RNA-seq from seeds of wild type *Arabidopsis*, FLOE1 deletion mutants, and mutants with the aspartic acid and serine (DS)-rich intrinsically disordered region deleted, which were dry, imbibed with plain water, or imbibed with salty water. The ∼6000 significant anchors found in this dataset include imbibition-induced cryptic splicing in AT2G36720, an acyl-CoA N-acyltransferase, with a splice junction from inside an intron to the 5’ boundary of the adjoining exon (Fig. 2C, SFig. S2C). Dry wild type seeds almost exclusively express the cryptic splice isoform, while water- and salt water-imbibed seeds express a mixture of isoforms, with ∼50-60% from the variant assigned to the canonical isoform. This condition-dependent splicing implies the possibility of yet-to-be discovered imbibition-dependent splicing regulatory programs in the seed and its role on the enzyme function.

In summary, SPLASH provides a highly efficient reference-free approach to detect multiple forms of sample-specific transcript diversification in plants. Here, we studied well-understood species with assembled genomes to evaluate this analytic framework by comparing it to well-established conventional genomic analysis. We foresee that more unbiased and high-throughput analysis of plant genomes will allow the plant genomics community to rapidly analyze genetic data from any plant, including those never before studied, without the tedious and time-consuming steps of assembly and alignment.

### Supplementary information

#### Total calls by SPLASH

To find additional examples of biologically interesting sequence variation with SPLASH, we examined unaligned variants whose expression varied by sample metadata such as experimental condition or developmental stage. First, we identified “unaligned” anchors, i.e. anchors for which one of the top two variants could not be aligned to the genome by STAR. Separately, we applied a generalized linear model (GLM) to all anchors to identify metadata-regulated anchors (Methods). Then, we looked for the anchors that were both “unaligned” to the genome as well as metadata-regulated, and investigated these resulting anchors starting with those having the largest effect size.

In a dataset of field-droughted sorghum^9^, there were 11,501 significant anchors (out of 544,476 total significant anchors, or 2.11%) for which one of the top two variants did not align to the genome. From the GLM output, there were 10,567 anchors with sample-dependent variation, corresponding to 3,829 unique genes (Supplementary Table 5). GO enrichment analysis on these regulated genes revealed 77 significantly enriched biological processes, with the top enriched being RNA splicing, via transesterification reactions with bulged adenosine as nucleophile (GO:0000377) and hydrogen peroxide catabolic process (GO:0042744). Of the regulated anchors in sorghum, 408 (3.86%) were also in the unaligned group.

One of the highest effect size anchors satisfying both conditions in the sorghum drought dataset had BLAST hits to fungal transcripts (see Supplementary Table 6 for full list). The most abundant variant had the best BLAST hits (with at least 96% coverage and 98.08% identity) that were uncharacterized genes from eight different fungal species in the genus *Alternaria*, which has been shown to be a human allergen^18^. For the second most abundant variant from this anchor, the best BLAST hits (all with 100% coverage and 94.44% identity) were annotated as hypothetical proteins in the species *Pseudogymnoascus verrucosus* and *Fusarium culmorum*, a wheat and sorghum pathogen^19^, or *Fusarium graminearum*, a species associated with toxicity in sorghum^20^ (Fig. 3A, Supplementary Fig. S3A). The most abundant variant differs between leaf and root samples, which might be explained by the presence of distinct fungal species on different tissues, or by tissue-regulated differential expression of related gene variants.

**Figure 3.**
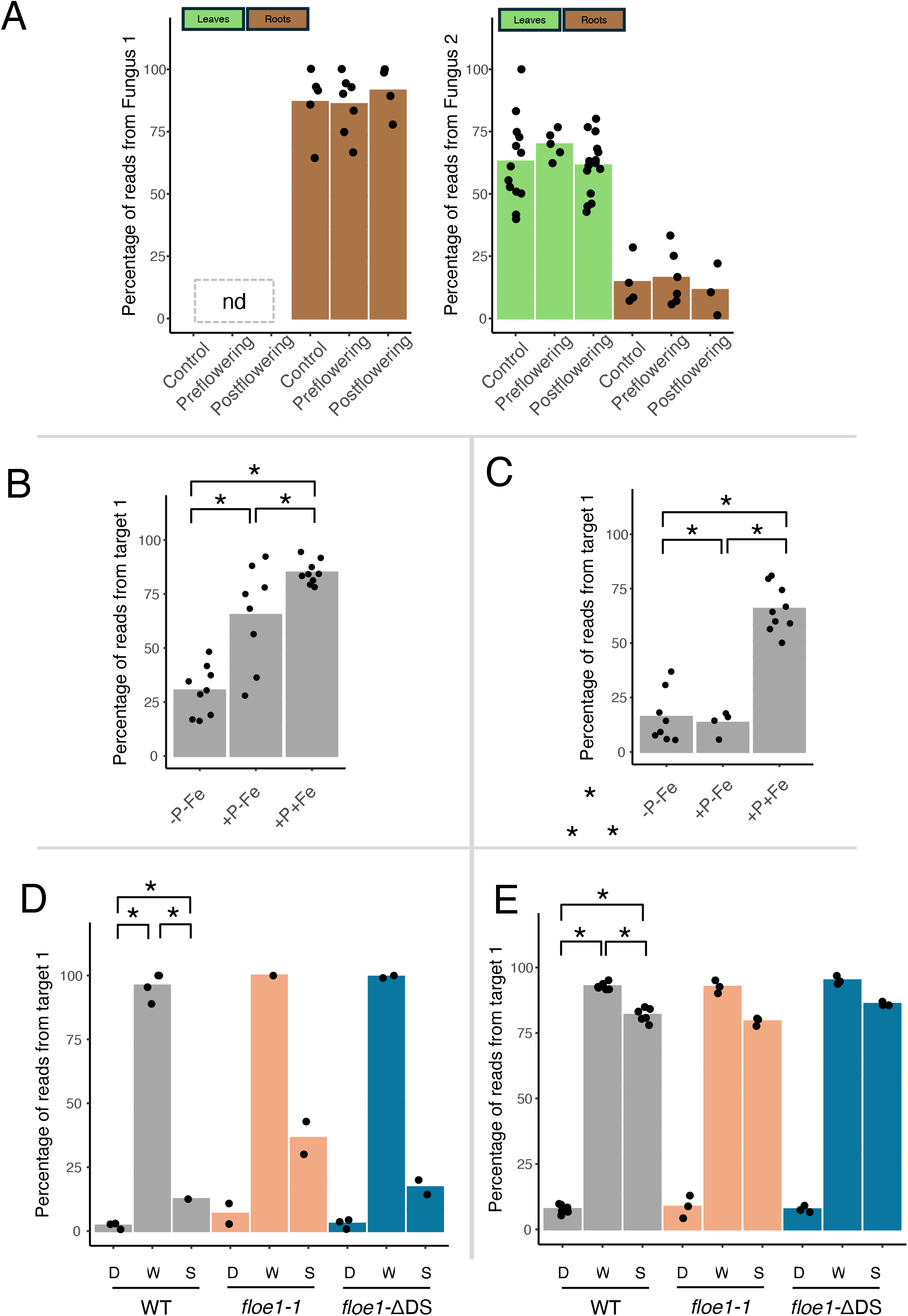
Proportion of reads within each sample from each of the top two variants, calculated as number of reads from that particular variant divided by total number of reads from the anchor. Bar height represents mean; individual datapoints are superimposed. Whether variation depends on experimental condition was tested in two different ways. For A, we used a generalized linear model to test whether the experimental condition/developmental stage could be predicted from the proportion of variants (Methods). For B-E, pairs of conditions were compared using a two-proportions z-test; asterisks indicate that the proportion of variant 1 reads is significantly different (p-value < 0.05) between experimental conditions (Methods). For the *Arabidopisis* FLOE1 dataset, only wild type (WT) samples were tested for significance. A: Sorghum drought dataset: the top three variants for this anchor BLAST to different fungal species. The most abundant variant had the best BLAST hits to fungal species in the genus *Alternaria*; the second most abundant variant had the best BLAST hits to species in the genera *Pseudogymnoascus* and *Fusarium* (see Supplementary Table 6 for full list). The abundance of each variant differs by tissue type (indicated by bar color). “Control” indicates samples with no drought stress; “preflowering” samples were droughted before the flowering stage; and “postflowering” samples were droughted after the flowering stage. N.d. means the anchor sequence was not found in the sequence data of samples in those conditions. This anchor was only found in 86 out of 198 total samples. B. *Arabidopsis* iron/phosphorus dataset: the first and second most abundant variants for this anchor align to homologous genes AT3G08720 (protein kinase 19) and AT3G08730 (protein-serine kinase 6) respectively; their relative expression varies by metadata condition. -P-Fe indicates the phosphorus and iron doubly deprived condition; +P-Fe is only iron deprivation; and +P+Fe indicates no deprivation. C. *Arabidopsis* iron/phosphorus dataset: the first and second most abundant variants for this anchor align to homologous genes AT1G62810 (copper amine oxidase 2) and AT3G43670 (copper amine oxidase 1) respectively; their relative expression varies by metadata condition. D: *Arabidopsis* FLOE1 dataset: the first and second most abundant variants for this anchor align to homologous squalene monooxygenase genes, AT5G24160 and AT5G24150; their relative expression varies by imbibition condition. WT (wild type), *floe1-1* (FLOE1 deletion mutant), and *floe1-ΔDS* (FLOE1 mutant with disordered region deleted) indicate the different seed genotypes; D (dry), W (wet), and S (salt) indicate the imbibition conditions for the seeds. Note: this anchor was only found in 28 out of 36 samples. E. *Arabidopsis* FLOE1 dataset: the first and second most abundant variants for this anchor align to homologous ERF/AP2 transcription factors, AT1G78080 (RAP2.4) and AT1G22190 (RAP2.4D); their relative expression varies by imbibition condition.

In the maize pollen dataset^10^ described in the main text, there were 551,639 anchors (out of 8,989,404 total significant anchors, or 6.14%) for which one of the top two variants did not align to the maize genome. Some of these unaligned variants can be explained by the mismatch between the maize hybrids used to generate the data (A188xB73) and the maize genome used for alignment (B73v5). From the GLM output, there were 190 anchors, corresponding to 78 unique transcripts/genes, with pollen-stage-specific expression (Supplementary Table 5). No biological processes were significantly enriched, suggesting that this phenomenon is general across biological processes and functions. Alternatively, the lack of statistical significance could be due to the small number of genes tested. Five (2.63%) of the regulated anchors were also in the unaligned group (Supplementary Table 6).

In the *Arabidopsis* iron/phosphorus deprivation dataset^12^ described in the main text, there were 134 anchors (out of 33,650 significant anchors, or 0.4%) for which one of the top two variants did not align. From the GLM output, there were 77 anchors with experimental condition-dependent variation, corresponding to 52 unique genes (Supplementary Table 5). There were no significantly enriched biological processes, perhaps due to the small number of genes tested. There was no overlap between the regulated anchors (as detected by the GLM approach) and the unaligned anchors.

In the *Arabidopsis* FLOE1 dataset^14^ described in the main text, there were 651 significant anchors (out of 224,642 total significant anchors, or 0.29%). From the GLM output, there were 6,470 sequences with condition- or genotype-specific expression, corresponding to 1,324 unique genes (Supplementary Table 5). GO enrichment analysis on the 651 significant condition-regulated genes for which one of the top two variants did not align produced 133 significantly enriched biological processes. The most enriched processes were glutamate biosynthetic process (GO:0006537), regulation of developmental vegetative growth (GO:1905613), positive regulation of chlorophyll biosynthetic process (GO:1902326) and mRNA splice site selection (GO:0006376). When we intersected these groups, there were 41 anchors in common (0.63% of condition-dependent anchors; see Supplementary Table 6).

There were also many significant anchors for which both of the top two variants could not be aligned to the genome (712 sequences in the *Arabidopsis* iron/phosphorus dataset; 46,932 in the *Arabidopsis* FLOE1 data; 131,618 in the sorghum data; and 1,099,637 in the maize data). For these anchors, the lists of sequences were so long that we could not query the sequences by BLAST in an efficient way; for this reason, we chose to investigate the category of anchors for which one of the top two variants did not align (as described above). Since we only investigated a small subset of anchors in depth, we expect there are many more examples of biologically important unaligned variants beyond the ones described in our study.

#### Additional discoveries from pairwise tests of metadata dependence

The GLM approach described above is conservative because condition-dependent variation may not be apparent from only considering the top two variants. To supplement the GLM approach, we used a second approach: we selected the top 500 anchors with the largest effect sizes in each dataset, and then tested whether the proportion of the top variant was significantly different between metadata groupings such as experimental condition or developmental stage (Methods).

In the *Arabidopsis* iron/phosphorus deprivation dataset^12^, we discovered multiple cases of differentially regulated homolog expression. For homologous genes AT3G08720 (protein kinase 19) and AT3G08730 (protein-serine kinase 6), relative expression of the homologs depended on phosphorus availability: in phosphorus-deficient samples, ∼70% of reads with this anchor correspond to expression of AT3G08730, while in samples grown in the presence of phosphorus, ∼70% of the reads with this anchor originate from AT3G08720 (Fig. 3B, Supplementary Fig. S3B). Another example is Copper Amine Oxidase: in iron-deficient samples, ∼90% of reads with this anchor correspond to expression of AT1G62810 (copper amine oxidase 2), while for samples grown in the presence of iron, ∼65% of reads with this anchor originate from AT3G43670 (copper amine oxidase 1) (Fig. 3C, Supplementary Fig. S3C).

We also discovered condition-dependent homolog expression in the *Arabidopsis* FLOE1 study^14^. For example, squalene monooxygenase genes, AT5G24160 and AT5G24150, are expressed differentially depending on imbibition conditions: dry and salt-water conditions induce the expression of AT5G24160, while imbibition with plain water results in expression of AT5G24150 (Fig. 3D, Supplementary Fig. S3D). For two homologs in the ERF/AP2 transcription factor family, AT1G78080 (RAP2.4) and AT1G22190 (RAP2.4D), the relative expression varies by whether the samples received any water. All dry samples express only AT1G22190, while imbibed samples express either AT1G78080 or a mix of both (Fig. 3E, Supplementary Fig. S3E).

Additionally, in the *Arabidopsis* FLOE1 dataset, we discovered regulated alternative splicing in the gene AT5G65080 (MAF5), which has been implicated in flowering timing in response to cold^22^. Samples from the dry condition tend to have the final intron spliced out, whereas salt and normal imbibition samples tend to include the final intron sequence.

#### Additional discoveries of cryptic splicing

Since SPLASH does not depend on using metadata, it can be used to discover variation that would not be found when looking for differences between experimental groups. For example, some samples in a dataset may express unannotated splicing isoforms, and SPLASH can discover these novel isoforms even if their expression did not differ by experimental condition. To find examples of splicing that were called significant by SPLASH, we aligned the significant anchors to the genome, and selected anchors with at least one splice junction among the associated variants. From this list, we prioritized cases where the variable sequences are the most dissimilar (Methods).

In the sorghum drought dataset, we detected unannotated splice junctions in genes SORBI_3002G381700 and SORBI_3003G250500 that encode hypothetical proteins, revealing splicing regulation in these poorly studied genes.

In the maize pollen dataset, we revealed unannotated splicing in the hypothetical protein Zm00001eb346620. In the *Arabidopsis* iron/phosphorus deprivation dataset, we found anchors where the different variants correspond to unspliced or spliced versions of the transcripts, for example in genes AT3G50480 (a homologue of RPW8, which is a disease resistance gene), AT2G47060 (cytosolic ABA receptor kinase 3), and pseudogene AT1G79245.

For an anchor in the *Arabidopsis* FLOE1 dataset, the associated variant sequences have a spliced alignment (aligning in two separate parts) upstream of the gene AT5G01530 into the 5’ untranslated region (UTR) of the gene. AT5G01530, also known as LHCB4.1, is part of the light-harvesting complex^23^. Splicing upstream of the annotated 5’UTR region could indicate an incomplete gene annotation, with splicing occurring within the 5’UTR^24^. However, these reads could also be explained by the presence of an upstream open reading frame (uORF), which could have strong regulatory effects such as translation inhibition^25^. In either case, regulated splicing upstream of the LHCB4.1 locus is likely to affect the resulting abundance of the LHCB4.1 protein. We also found cases of splice junctions into introns, for example in genes AT1G60900 (a putative U2A65 splicing factor involved in flowering regulation) and AT1G80570 (an RNI-like superfamily protein).

#### Validation of previously unannotated splice junctions

In the *Arabidopsis* iron/phosphorus deprivation dataset, we also found instances of unannotated alternative splicing with a low effect size, indicating that there is not a large difference in which isoforms are expressed in different samples (effect size < 0.053). AT1G13609, a defensin-like (DEFL) protein known to be regulated by iron deficiency has unannotated splice junctions from within the final exon to downstream of the annotated 3’ UTR end (Supplementary Fig. S4A). We validated this unexpected splicing with amplicon sequencing of the region from PCR-amplified cDNA (Methods). The predicted alternative splicing of AT1G79245 was similarly validated (Supplementary Fig. S4B-E; Methods).

In summary, by bypassing reference alignment, SPLASH reveals complex regulation of sequence diversification mechanisms, including alternative splicing and differential homolog expression. SPLASH represents a critical step forward in plant genomics – here applied to RNA, but also applicable to DNA – that enables rapid, precise discovery of genomic regulation and functional prioritization without an alignment approach.

## Methods

### SPLASH runs

The implementation of SPLASH used in this paper is described in^8^. Specifically, SPLASH version 1.9.0 (https://github.com/refresh-bio/SPLASH/tree/archive/1.9.0) was run on each dataset. When paired end reads were available, only R1 reads (i.e., the first read from each pair of reads) were used as input for SPLASH. This was done because SPLASH does not currently have a way to process paired end reads. For the sorghum drought dataset, only samples from the BTX642 genotype were input for SPLASH. For the other datasets, all samples were used as input for SPLASH.

The same following parameters were used for each SPLASH run: n_bins = 128; max_pval_rand_init_alt_max_for_Cjs = 0.1; anchor_len = 27; target_len = 27; gap_len = 0; poly_ACGT_len = 8; anchor_unique_targets_threshold = 1; anchor_count_threshold = 50; anchor_samples_threshold = 1; anchor_sample_counts_threshold = 5; n_most_freq_targets = 10; generate_alt_max_cf_no_tires = 10; altMaximize_iters = 50; train_fraction = 0.25; kmc_use_RAM_only_mode = True; calculate_stats = True; without_SVD = True; with_effect_size_cts = False; enable_pvals_correction = True; fdr_threshold = 0.05.

An example of running the command looks like the following “docker run -v ‘pwd’:/home/ ubuntu ghcr.io/refresh-bio/splash:1.9.0 splash --n_threads_stage_1 3 --n_threads_stage_2 8 -- n_bins 128 --gap_len 0 --calculate_stats --dump_Cjs --n_most_freq_targets 10 -- pvals_correction_col_name pval_rand_init_alt_max --enable_pvals_correction --without_SVD -- clean_up --kmc_use_RAM_only_mode input.txt”. Where “--n_threads_stage_1” and “-- n_threads_stage_2” determine the number of threads used in each stage based on the memory usage of the species. “Input.txt” is a space-delimited file that maps the input samples’ names to their file paths.

The SPLASH output consists of significant sequences (“anchors”) and the associated diversified sequences (“targets”). SPLASH also automatically outputs some summary statistics including effect size, number of unique targets per anchor, average Hamming distance between each target and the top target, etc.

### Local assembly of anchors

Local assemblies based on each anchor were generated from reads using the method described in ^26^.

### Filtering out molecular biology artifacts

To remove false positive anchors that originate from sequences present in molecular biology tools, such as sequencing adapters, we used Bowtie2 to align anchors against indices generated from the UniVec database (obtained from ftp://ftp.ncbi.nlm.nih.gov/pub/UniVec/) and a set of Illumina adapters. If an anchor aligned to either database, it was discarded as an artifact.

### Alignment of anchor/target sequence to genome

All reference genome FASTA and BED files were downloaded from Ensembl Plants. For Arabidopsis, we aligned to the TAIR10 assembly (https://ftp.ensemblgenomes.ebi.ac.uk/pub/plants/release-56/fasta/arabidopsis_thaliana/); for sorghum, we aligned to the NCBIv3 assembly (https://ftp.ensemblgenomes.ebi.ac.uk/pub/plants/release-56/fasta/sorghum_bicolor/); and for maize, we aligned to the B73 AGPv5 assembly (zeaMay_b73_v5; https://ftp.ensemblgenomes.ebi.ac.uk/pub/plants/release-56/fasta/zea_mays/).

The alignment and gene name assignment approach was adapted from^15^. For each significant anchor in each dataset, we concatenated the anchor and target sequence for up to ten targets reported by SPLASH, and saved all these concatenated sequences as FASTA files. Then, we aligned each concatenated anchor/target sequence to the respective plant genome using STAR version 2.7.5a. This is STAR command that was used for alignment: STAR --runThreadN 4 --genomeDir <star_index> --readFilesIn <concatenated_anchor_target_fasta_file> --outFileNamePrefix <output_folder> --twopassMode Basic --alignIntronMax 1000000 --chimJunctionOverhangMin 10 --chimSegmentReadGapMax 0 --chimOutJunctionFormat 1 --chimSegmentMin 12 --chimScoreJunctionNonGTAG -4 -- chimNonchimScoreDropMin 10 --outSAMtype SAM --chimOutType SeparateSAMold -- outSAMunmapped None --clip3pAdapterSeq AAAAAAAAA --outSAMattributes NH HI AS nM NM

Gene names were assigned by extracting exon positions from the STAR BAM output and applying the bedtools function “intersect” to the exon positions as well as a reference BED file of gene and exon boundaries.

To assign anchors to genes/transcripts, the anchors were aligned using Bowtie2 to the respective reference genome/transcriptome.

### Alignment of anchor to database of repetitive sequences

We downloaded databases of repetitive elements in *Arabidopsis thaliana, Sorghum bicolor*, and *Zea mays* from the Plant Repeat Database (Plantrep) at http://www.plantrep.cn/^16^. We then used Bowtie2 to create indices from these databases, and then aligned anchor sequences to these indices.

### Querying BLAST

Anchor and target sequences were concatenated and saved as FASTA files. We submitted each FASTA file as a query to BLAST using the following command: blastn -outfmt 6 qseqid sseqid pident length mismatch gapopen qstart qend sstart send evalue bitscore sseqid sgi sacc slen staxids stitle -query <fasta_path> -remote -db nt –out <blast_output> -evalue 0.2 -task blastn -dust no -word_size 24 -reward 1 -penalty -3 -max_target_seqs 20

The BLAST hits were sorted by increasing E-value, so the “top” hit is the one with the smallest E-value.

For A118 and B73 allele confirmation, MaizeGDB BLAST (https://maizegdb.org/popcorn/search/sequence_search/home.php?a=BLAST_UI) were used to perform anchors/targets BLAST against A188 and B73 genome sequences.

### Generalized linear model (GLM) to detect metadata-dependent target usage

To automatically detect anchors whose target usage (variant expression) varies depending on the metadata grouping, we used the R package “glmnet” to run a GLMnet Lasso multinomial regression. If the counts from the top two targets of an anchor can predict metadata category, i.e. the largest GLM coefficient is greater than one, then the anchor is classified as metadata-dependent.

We began by identifying “unaligned” anchors, i.e. anchors for which one of the top two targets had a concatenated anchor/target sequence that could not be aligned to the genome by STAR. We also applied the GLM to all anchors to identify metadata-regulated anchors. Finally, we intersected the group of “unaligned” anchors with the group of metadata-regulated anchors, and investigated the resulting anchors starting with those having the largest effect size.

### Filtering high-effect anchors in *Arabidopsis* FLOE1 and maize pollen

To find examples of anchors with condition-dependent target usage, we initially ranked anchors by decreasing effect size. However, for the *Arabidopsis* FLOE1 and maize pollen datasets, most of the anchors with the highest effect sizes were only present in low numbers of reads. To keep only anchors with robust representation in the raw reads, we required that at least seven of concatenated anchor/target sequences from the top ten targets aligned to the genome. We then filtered for only anchors present in at least 500 reads. Finally, we ranked anchors by decreasing effect size and selected the top 500 anchors to inspect for condition-dependent target usage.

### Testing whether the fraction of the top target is significantly different by metadata category

To find examples of anchors with condition-dependent target usage from the lists of high-effect size anchors, we began by visually inspecting plots showing expression of each target as grouped by metadata category. For anchors that appeared to have metadata-dependent target usage, we applied a two-proportions z-test (using the function prop.test in R) comparing the fraction of reads from target 1 in pairs of metadata conditions.

### Plant Materials for independent validation

*Arabidopsis thaliana* Col-0 seeds were surface sterilized using 70% Ethanol for 10 minutes, then plated on 0.5x Murashige and Skoog (MS) plates (PhytoTechnologies Laboratories) (pH = 5.7). Seeds were stratified at 4C for 48hrs in the dark, then placed upright in growth chambers maintaining constant temperature of 23C, with 150μmol white light provided under long (16hr) day conditions. After 10 days of growth, seedlings (10-15) from individual plates were consolidated and immediately frozen and ground to a fine powder in liquid nitrogen before long-term storage at -80C.

### RNA extraction and cDNA library preparation

RNA was extracted from 100mg of frozen *Arabidopsis* tissues using the RNEasy Plant Mini Kit (Qiagen) per manufacturer recommendations. cDNA was synthesized from 2μg RNA using M-MLV Reverse Transcriptase (Thermo Scientific). DNA contamination was assessed by performing PCR amplification over a characterized intron junction and assessing intron retention (genomic DNA contamination) by agarose gel electrophoresis. Samples that produced a single band, of a size corresponding to properly spliced mRNA, were used for downstream analysis.

### PCR and amplicon sequencing

For validation of splicing in AT1G13609, the region of interest was amplified from cDNA using DreamTaq master mix (Thermo Scientific). Primers as denoted in SFig. 4A (5’ to 3’):

Forward primer: GCGTAATTATGTCAGTGTTATTGGC

Reverse primer: GCTTCTTCTCATCCAGTTTACAAGC

The resulting products were visualized on a 1.5% agarose gel, purified, and submitted to the NGS Amplicon-EZ service from Azenta Life Sciences.

For genes where alternative splice forms could be separated through electrophoresis (AT1G79245), individual bands were purified separately and submitted for Sanger sequencing from Sequetech Corporation.

Primers as denoted in SFig. 4B-D (5’ to 3’):

Primer_fw: GCCTGGAATCTGCACAAGTTG

Primer_rev: TTACTGAAGTTATCATGGGAAGCACT

## Data availability

All FASTQ files were downloaded from the Sequence Read Archive (SRA). The BioProject accession numbers for each dataset are: *Arabidopsis* FLOE1 - PRJNA704067 and PRJNA704079; *Arabidopsis* iron/phosphorus - PRJNA685167; sorghum drought - PRJNA527782; and maize pollen - PRJNA732658 and PRJNA734295.

## Author contributions

EM analyzed data generated from SPLASH, worked on experimental validation of SPLASH results, and contributed to writing the manuscript. EVS analyzed data generated from SPLASH, worked on experimental validation of SPLASH results, and contributed to writing the manuscript. MK contributed to development of the SPLASH method. BX analyzed data generated from SPLASH and contributed to writing the manuscript. SD contributed to development of the SPLASH method. SYR oversaw the project and contributed to writing the manuscript. JS oversaw the project, contributed to development of the SPLASH method, and contributed to writing the manuscript.

## Acknowledgements

We thank Roozbeh Dehghannasiri for a wrapper script to run STAR and for the GLM analysis script. We thank members of the Salzman and Rhee labs for helpful discussions. This work was supported in part by the National Science Foundation Graduate Research Fellowship Program (grant number DGE-1656518), a training grant from NIH Cellular and Molecular Training Grant (NIGMS, grant number 5T32GM007276), US National Science Foundation grants (MCB-1617020, IOS-1546838) (S.Y.R.), Water and Life Interface Institute (WALII) DBI (grant no. 2213983) (S.Y.R.), US Department of Energy, Office of Science, Office of Biological and Environmental Research, Genomic Science Program grant nos (DE-SC0018277, DE-SC0008769, DE-SC0020366, DE-SC0023160, and DE-SC0021286) (S.Y.R.). This work was done on the ancestral land of the Muwekma Ohlone Tribe, which was and continues to be of great importance to the Ohlone people.

---

**Supplementary Figure S1.**
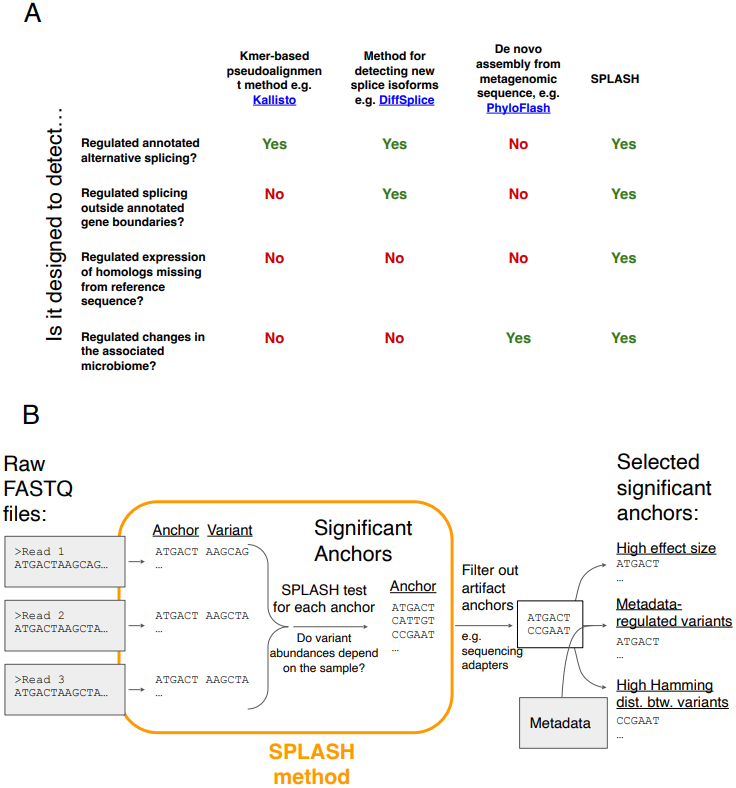
A. Comparison between SPLASH and other types of genomics analysis, showcasing the unique strengths of SPLASH. B: Overview of workflow for identifying anchors of interest. SPLASH takes in raw reads in FASTQ format and identifies “anchor” sequences (kmers) that precede different “variant” sequences. For each such anchor, SPLASH runs a statistical test to determine whether the abundance of each variant varies by sample. The significant anchors produced by SPLASH can then be prioritized by the user in various ways, such as selecting anchors with a large SPLASH effect size, selecting anchors where the variant usage differs by metadata condition, selecting anchors with a high Hamming distance between variants, and more. See Methods for details.

**Supplemental Figure 2.**
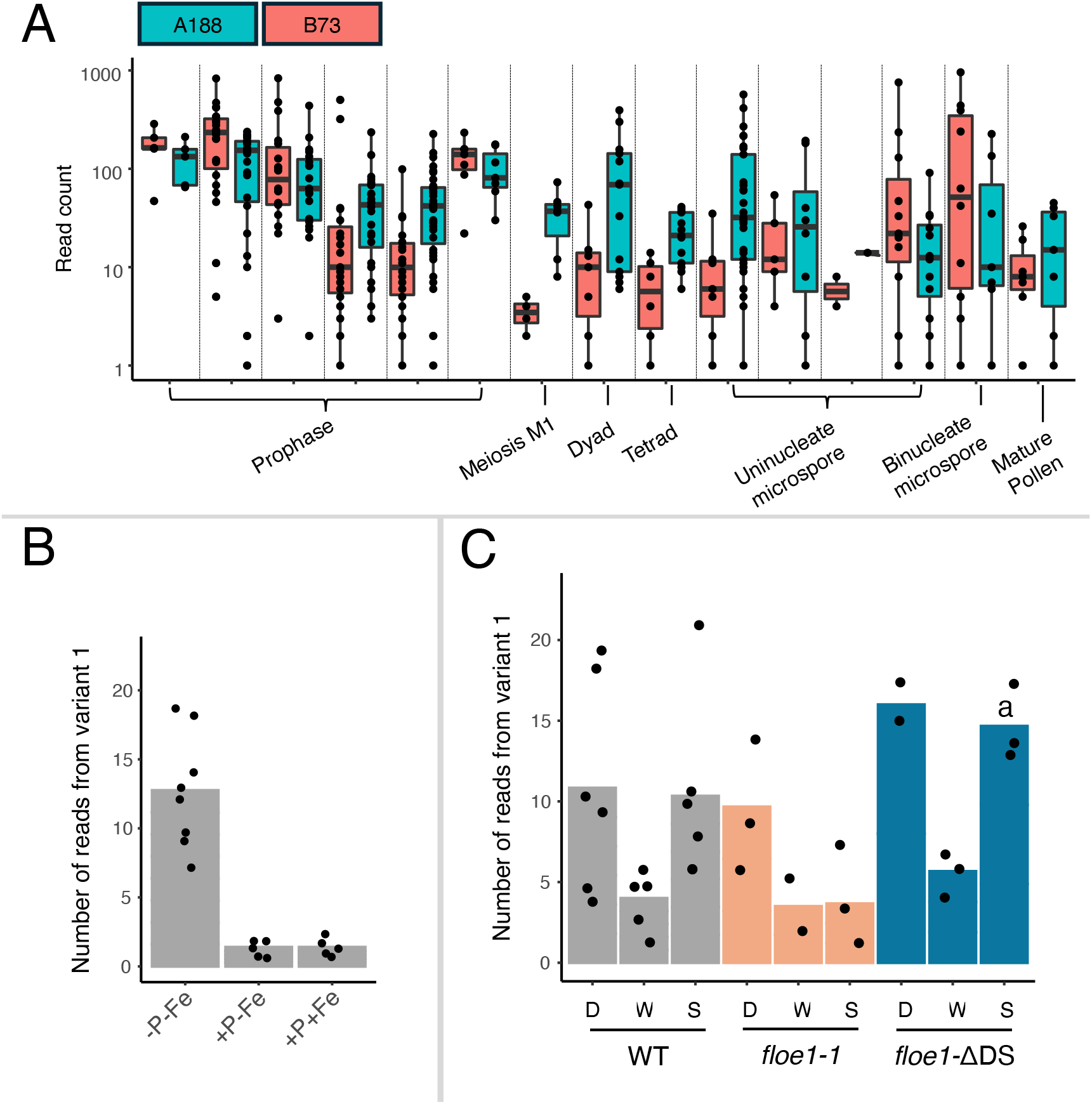
Number of raw reads within each sample from each of the top two variants. Bar height represents mean; individual datapoints are superimposed. A: Maize pollen dataset: the top two variants for this anchor align to alleles of Zm00001eb173470 from B73 and A188. The differential expression of these alleles varies by pollen stage. Note: this anchor was only found in 260 out of 642 samples. B: *Arabidopsis* iron/phosphorus deprivation dataset: variant 1 maps to a splice junction in AT1G74270 (ribosomal protein L35Ae), while variant 2 includes the intron. -P-Fe indicates the phosphorus and iron doubly deprived condition; +P-Fe is only iron deprivation; and +P+Fe indicates no deprivation. C: *Arabidopsis* FLOE1 dataset: variant 1 maps to an annotated splice junction between exons in AT2G36720, but variant 2 maps to a cryptic splicing event from inside an intron to an exon. WT (wild type), *floe1-1* (FLOE1 deletion mutant), and *floe1-ΔDS* (FLOE1 mutant with disordered region deleted) indicate the different seed genotypes; D (dry), W (wet), and S (salt) indicate the imbibition conditions for the seeds.

**Supplemental Figure 3.**
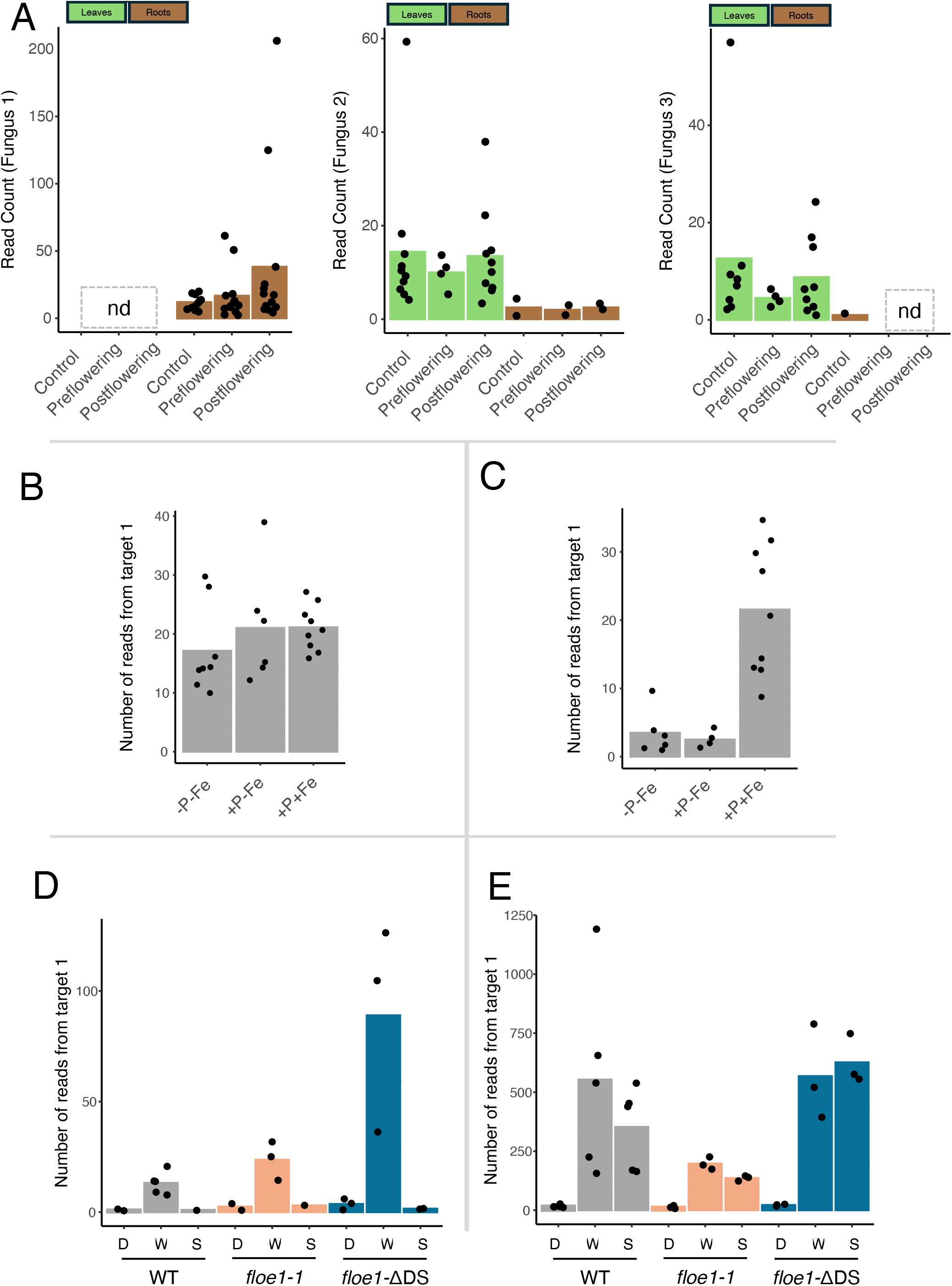
Number of raw reads within each sample from each of the top two variants. Bar height represents mean; individual datapoints are superimposed. A: Sorghum drought dataset: the top three variants for this anchor BLAST to different fungal species. The most abundant variant had the best BLAST hits to fungal species in the genus *Alternaria*; the second most abundant variant had the best BLAST hits to species in the genera *Pseudogymnoascus* and *Fusarium* (see Supplementary Table 6 for full list). The abundance of each variant differs by tissue type (indicated by bar color). “Control” indicates samples with no drought stress; “preflowering” samples were droughted before the flowering stage; and “postflowering” samples were droughted after the flowering stage. N.d. means the anchor sequence was not found in the sequence data of samples in those conditions. This anchor was only found in 86 out of 198 total samples. B. *Arabidopsis* iron/phosphorus dataset: the first and second most abundant variants for this anchor align to homologous genes AT3G08720 (protein kinase 19) and AT3G08730 (protein-serine kinase 6) respectively; their relative expression varies by metadata condition. -P-Fe indicates the phosphorus and iron doubly deprived condition; +P-Fe is only iron deprivation; and +P+Fe indicates no deprivation. C. *Arabidopsis* iron/phosphorus dataset: the first and second most abundant variants for this anchor align to homologous genes AT1G62810 (copper amine oxidase 2) and AT3G43670 (copper amine oxidase 1) respectively; their relative expression varies by metadata condition. D: *Arabidopsis* FLOE1 dataset: the first and second most abundant variants for this anchor align to homologous squalene monooxygenase genes, AT5G24160 and AT5G24150; their relative expression varies by imbibition condition. WT (wild type), *floe1-1* (FLOE1 deletion mutant), and *floe1-ΔDS* (FLOE1 mutant with disordered region deleted) indicate the different seed genotypes; D (dry), W (wet), and S (salt) indicate the imbibition conditions for the seeds. Note: this anchor was only found in 28 out of 36 samples. E. *Arabidopsis* FLOE1 dataset: the first and second most abundant variants for this anchor align to homologous ERF/AP2 transcription factors, AT1G78080 (RAP2.4) and AT1G22190 (RAP2.4D); their relative expression varies by imbibition condition.

**Supplemental Figure 4.**
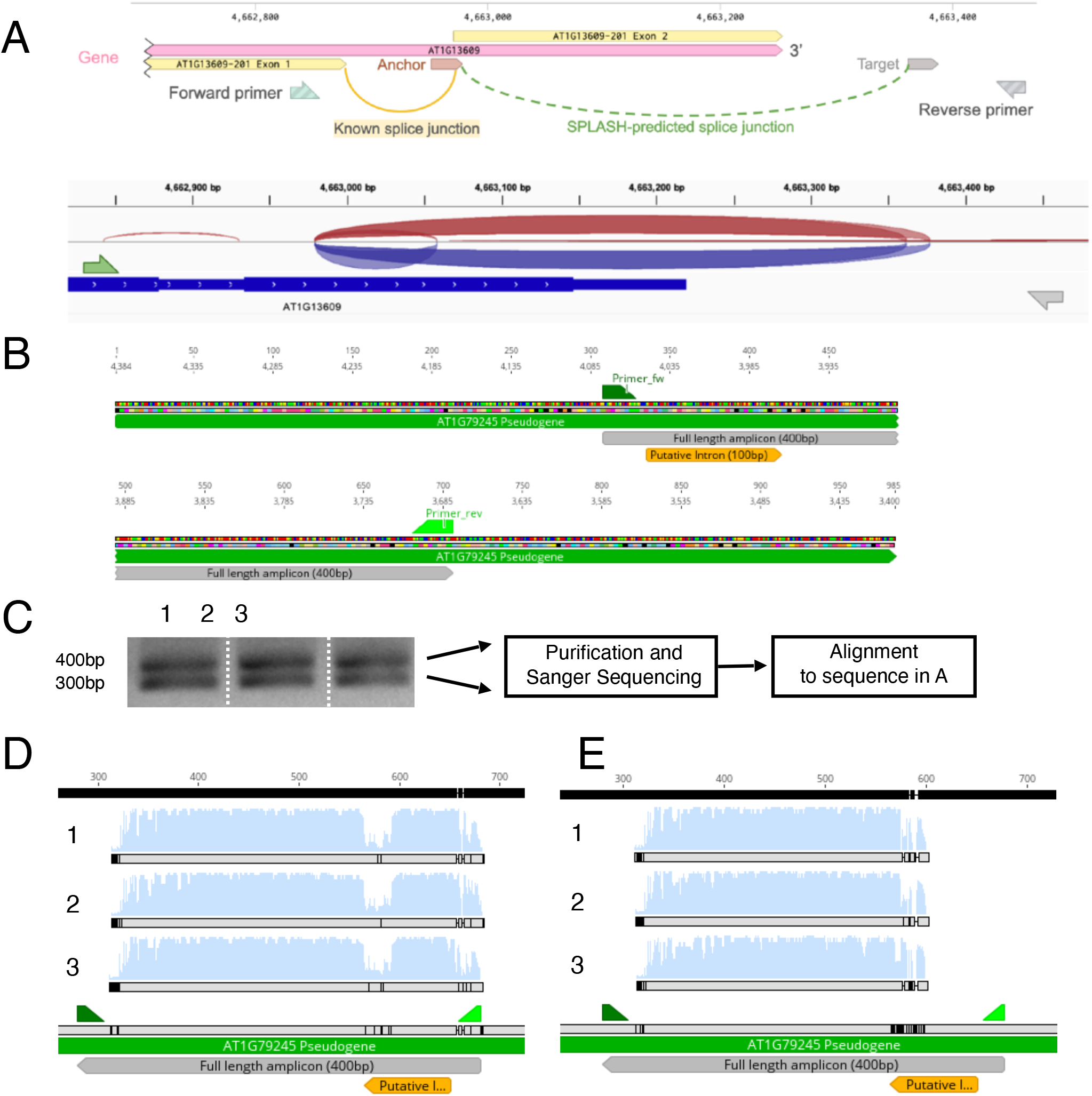
A: Validation of alternative splicing in AT1G13609. Above: SPLASH predicts a previously unannotated splice junction (dotted green line). Primers were designed surrounding this predicted junction. Below: The resulting PCR product was purified and sequenced via NGS Amplicon Sequencing (see Methods for details). The red and blue arcs represent reads with spliced alignments, confirming the presence of an unannotated splice junction spanning from the final exon of the gene to the 3’ UTR region. B-D: Validation of alternative splicing in the AT1G79245 pseudogene. (B) SPLASH annotates a 100bp putative intron in the *Arabidopsis* pseudogene AT1G79245. Nucleotide coordinates are provided above the sequence as grey numbers. The top row indicates positions within the represented 1kb fragment, starting at position 1. The bottom row indicates positions within the AT1G79245 pseudogene. AT1G79245 is encoded on the (-) strand of the *Arabidopsis* genome. Coordinates are provided according to the (+) strand of the genome, hence the descending position number. Predicted binding sites for Primer_fw and Primer_rev, calculated using the Primer_bind algorithm from Geneious, are shown as green trapezoids above the sequence. The predicted full length amplicon, and the position of the putative intron predicted by SPLASH are shown, with predicted 400bp (Intron retained) and 300bp (Intron spliced out) PCR products following amplification with Primer_fw and Primer_rev. C: Banding pattern on a 1% Agarose Gel following PCR amplification with Primer_fw and Primer_rev, from cDNA derived from 3 independent Col-0 seedling pools. Band sizes were estimated based on comparison to the GeneRuler 100bp ladder. A schematic of downstream analysis is shown in to the right of the visualized gel. Sanger Sequencing was performed using Primer_fw to prime the reaction. D-E: Alignments of Sanger Sequencing Results using DNA purified from the top (D) and bottom (E) bands shown in C. Per-nucleotide quality for samples 1-3 is shown as blue bars above the consensus sequence alignment. Alignment fidelity is shown as a grey bar for each sample, with dark lines or gaps indicating a failure to align, and light grey indicating a 100% sequence identity between samples.

## Notes

### Competing Interest Statement

The authors have declared no competing interest.

